# High Resolution Outdoor Videography of Insects Using Fast Lock-On Tracking

**DOI:** 10.1101/2023.12.20.572558

**Authors:** T. Thang Vo-Doan, Victor V. Titov, Michael J.M. Harrap, Stephan Lochner, Andrew D. Straw

## Abstract

Insects have significant global impacts on ecology, economy, and health and yet our understanding of their behavior remains limited. Bees, for example, use vision and a tiny brain to find flowers and return home, but understanding how they perform these impressive tasks has been hampered by limitations in recording technology. Here we present Fast Lock-On (FLO) tracking. This method moves an image sensor to remain focused on a retroreflective marker affixed to an insect. Using paraxial infrared illumination, simple image processing can localize the sensor location of the insect in a few milliseconds. When coupled with a feedback system to steer a high magnification optical system to remain focused on the insect, a high spatial-temporal resolution trajectory can be gathered over a large region. As the basis for several robotic systems, we show FLO is a versatile idea which can be employed in combination with other components. We demonstrate that the optical path can be split and used for recording high-speed video. Furthermore, by flying a FLO system on a quadcopter drone, we track a flying honey bee and anticipate tracking insects in the wild over kilometer scales. Such systems have the capability of providing higher resolution information about insects behaving in natural environments and as such will be helpful in revealing the biomechanical and neuroethological mechanisms used by insects in natural settings.

**One-Sentence Summary:** Fast Lock-On tracking enables recording trajectories and high-speed videos of insects behaving over large areas in the wild.

## INTRODUCTION

Insects, particularly pollinators like bees, are crucial for ecosystem stability and food security through their role in pollination, which is essential for the reproduction of many plants, including numerous crops, and the sustainability of our global food supply [1]. Like other insect groups, many wild bee species are currently suffering dramatic losses in abundance, likely due to contributions from several factors [2]. Studying insect behavior is an important step in revealing the drivers of these losses and their mechanisms of action. Classically, such behavioral research has been done by direct observation. This has revealed, for example, the waggle dance by which forager bees communicate to naïve nestmates the direction and distance to food sources [3]. More recently, radar tracking of insects has been developed [4, 5] and, for example, used to study the development of kilometer-scale flight trajectories over the lifetime of a bee [6]. Despite unique capabilities to track over landscape-scales, radar suffers from limited resolution in the spatial and temporal domains (Fig. 1A). To obtain more numerous and higher resolution data, researchers have built sophisticated laboratory setups with high resolution cameras in a smaller volume. These have revealed, for example, the biomechanical and sensory basis for decisions made in flight [7, 8] but, as they are in the lab, are not able to study insects in their natural environment where many additional factors may influence performance.

**Fig. 1.**
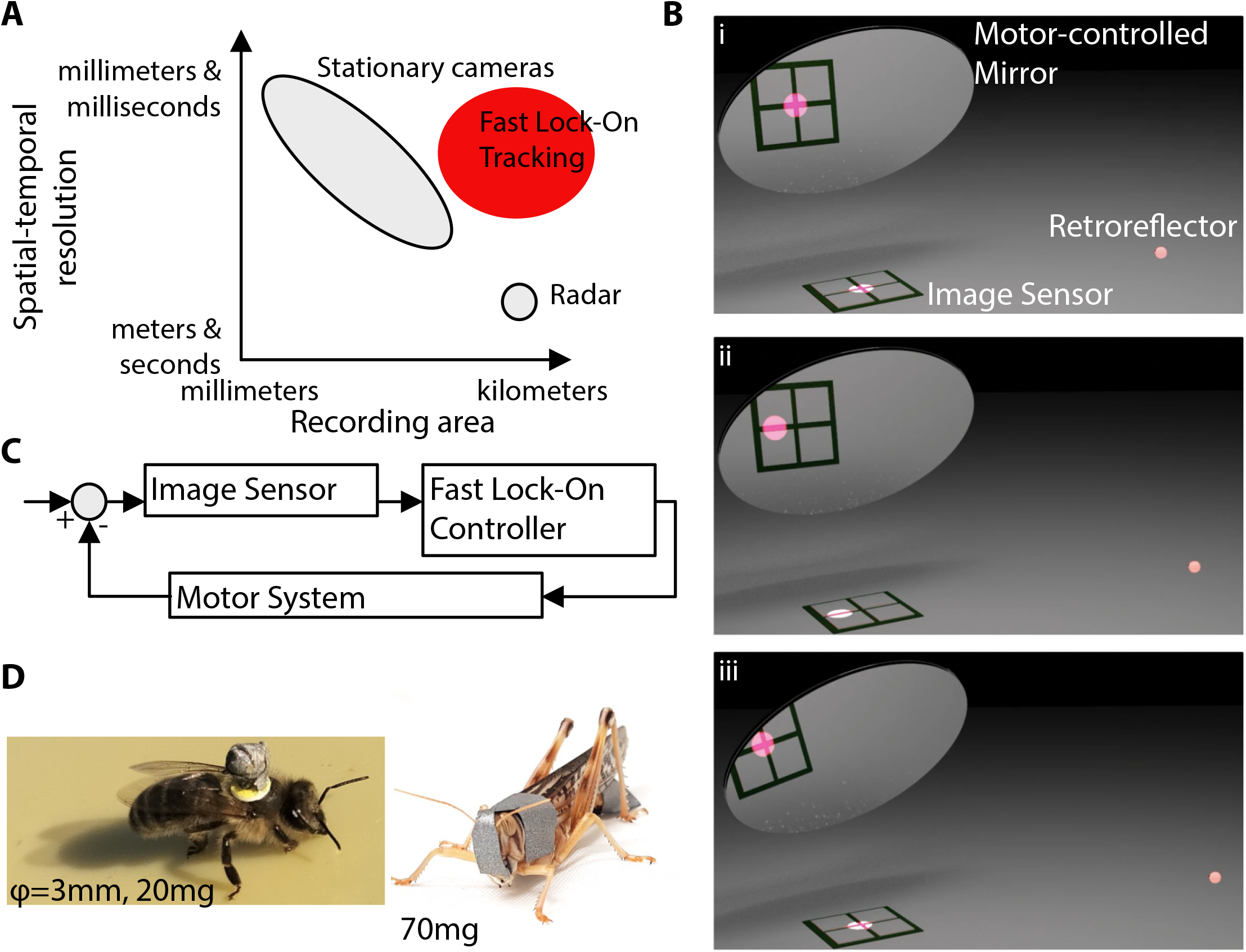
Fast lock-on tracking bypasses tradeoffs of other insect tracking techniques. (**A**) Conceptual view of tracking systems based on stationary cameras, radar, and Fast Lock-On (FLO) Tracking. Moving cameras allow expanding the recording area beyond stationary cameras, but without a Fast Lock-On system it has not been possible to follow insects with a moving camera due to their small size and speed. (**B**) At the starting configuration (i), the system is aimed such that the image of the insect-mounted reflector is focused in the center of the image sensor. Note that illumination and focusing optics are omitted from the illustration. When the insect moves, the reflected spot is initially offset from the sensor center (ii). Based on image processing and motor action, the system automatically adjusts mirror position to re-center the spot with low latency (iii). (**C**) Schematic block diagram showing that the input to the image sensor is the angular difference between the insect angle and the motor-controlled optical path angle. (**D**) Examples of insects carrying representative retroreflective markers tracked with FLO systems include the honey bee *Apis mellifera* (marker diameter 3 mm, mass 20 mg) and the locust *Schistocerca gregaria* (marker mass 70 mg).

Videography of flying insects has been an important methodology to quantify their behavior but a stationary camera is subject to a fundamental tradeoff imposed by the fixed number of pixels [9]. If a larger volume is to be filmed, the pixels must capture a wider area and hence have lower angular resolution. While adding more pixels is possible to sidestep the issue, the image of a moving subject also moves on the sensor, thus creating motion blur. A standard method to overcome these challenges has been simply to reduce relative motion between the subject and the camera by using a high magnification objective and continuously adjusting its aim [10]. In the case of flying insects, however, these are difficult to track with the naked eye and nearly impossible to keep a camera aimed and focused without automation. Use of low-latency optical processing has been used for decades in microscopy to track bacteria, *Paramecia* and roundworms. In these systems, an optical position signal is used to adjust the microscope stage such that the subject remains at the system’s focus [11, 12, 13]. For an animal or robot moving at larger spatial scales, however, another solution is necessary. For inanimate objects such as sports balls or drones and for larger animals such as birds, high-magnification optics have been combined with high-speed image analysis to enable tracking and videography [14, 15, 16]. Tracking insects with such systems, however, remains challenging. Simplifying the complexity of the tracking task to achieve the necessary low latency is essential, and one solution is to use simplified indoor backgrounds [17]. Another solution is to translate a frame carrying multiple cameras around a flying insect using multiple winches comprising a cable robot within a specially equipped space [18].

Affixing an optically distinct marker, such as one that is much brighter than the background or has a unique color, could simplify the image processing task and hence make it robust and fast. One choice would be to use a fluorescent or luminescent marker, which works in sunlight [19]. However, the requirement of sunlight is potentially limiting, for example to study the nocturnal insects and the effects of artificial light at night. Even worse, however, is that omnidirectional emission of fluorescence and luminescence greatly reduces the intensity, and hence maximum tracking range, of returned light. On the other hand, retroreflectors return most incoming energy to within a small angle of the incident direction, and may be suitable as markers when combined with paraxial illumination such as an LED near the camera lens. Retroreflective markers are widely used in commercial (human) motion-capture systems, and have already been used in a variety of insect tracking tasks [7, 20, 21, 22]. Our own preliminary work was promising [23] and here we report a full follow-up.

Inspired by these earlier works which implemented what we here call Fast Lock-On (FLO) tracking, we sought to apply FLO to the task of tracking insects flying outdoors, which remains an incompletely solved yet important problem.

## RESULTS

### Implementation of Fast Lock-On tracking

The basic principle of Fast Lock-On (FLO) tracking is to use feedback from an optical sensor to steer its optical axis, e.g. by tilting a mirror, in a manner that minimizes deviation of the target image from the center of the sensor (Fig. 1B). The time history of the optical axis position and sensor output can be used to compute the tracked object’s trajectory. In our case, we pan and tilt the optical axis, and this allows us to reconstruct the flight path of an insect in angular terms as measured from the FLO system. Low latency from sensor input to motor output will improve system performance and enable higher magnification optics, which in term can lead to further performance gains. From a control perspective, FLO is a closed-loop design in which the displacement of the target image from the sensor center is an error angle between the insect angle and the angular position of the optical path under control. The FLO controller sends motor commands to minimize this error and receives feedback through the optical sensor (Fig. 1C). To track insects using FLO systems, we incorporate infrared illumination near the optical axis of the optical sensor and mark the insect with a small, lightweight retroreflective marker (Fig. 1D).

A simple FLO system can be realized with a handful of inexpensive parts, namely a low-latency (on the order of 10 milliseconds) digital camera, a pan-tilt motor system, and a computer (Fig. 2, Supplemental Materials). Despite being relatively simple, this system is useful as a starting point for more sophisticated hardware and experimenting with different image processing and control algorithms. To reduce latency of image processing, we implemented only a very simple bright point detector. On the control side, while it is possible to implement FLO tracking with an internal model estimating only the angular error of the target with respect to the moving optical axis and updating motor commands accordingly, we found that using a Kalman filter to estimate both angular position and velocity of the target with respect to a fixed reference frame improved tracking robustness as the system was then able to maintain tracking during occlusions.

**Fig. 2.**
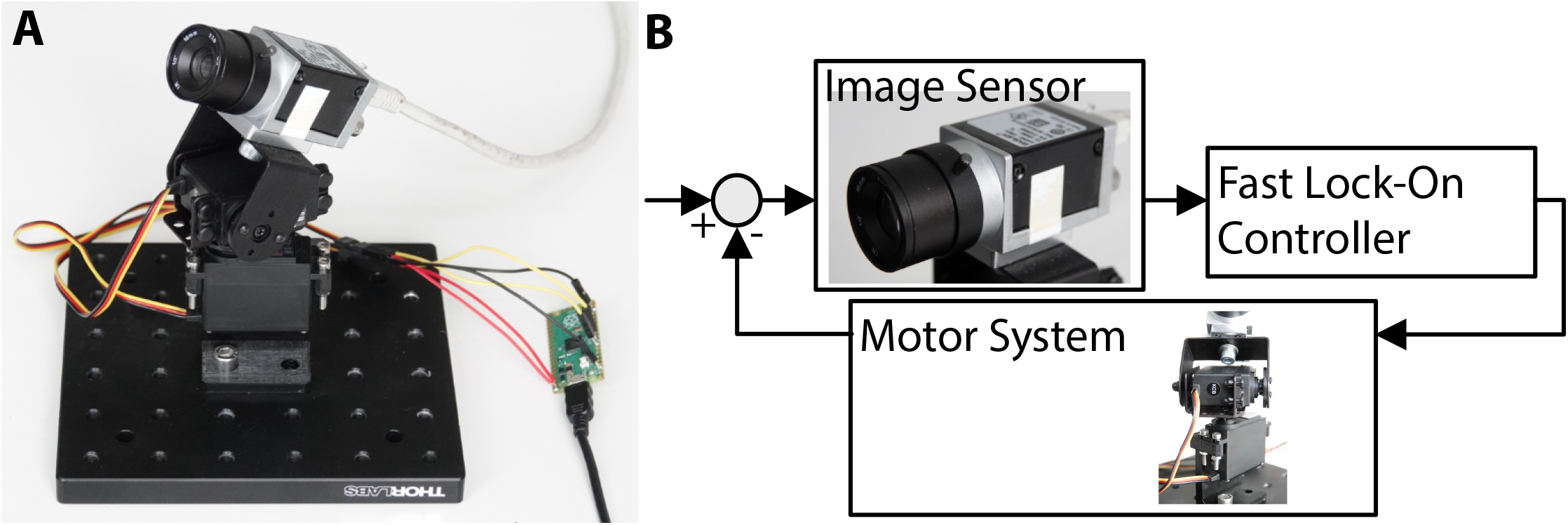
Simple fast lock-on tracking system. (**A**) For prototyping algorithms and software, a simple pan-tilt system can be built from commercial off the shelf parts. (**B**) The system architecture allows gaining experience in Fast Lock-On tracking.

Because FLO tracking views the target through the optical path under control, calibration is simplified compared to a system in which a separate camera system is used to steer the optical path for a high-magnification sensor [24, 25]. A low latency closed loop performance contributes to improved robustness not only through simply increasing the frequency of disturbances which can be compensated, but also by minimizing the maximum displacement over time of a moving target and thus reducing the difficulty for a tracking system to distinguish which of multiple potential targets corresponds to the object being tracked. With the use of beam splitters or selective transmission mirrors, “copies” of the optical path can be built and imaged on cameras with different characteristics than the tracking camera.

### High-speed, high-resolution videos of insects flying outdoors

To make high resolution videos of insects flying outdoors, several challenges remain. One challenge is that direct or reflected sunlight can be detected by our simple image processing instead of the insect-attached marker. Therefore, we sought to minimize the recorded brightness of sunlight and maximize the intensity of the retroreflective target image by confining our illumination and detection to the same, narrowly defined temporal and spectral windows. This maximizes the signal recovered from the illumination power but minimizes the noise from sunlight. We used high-power pulsed illumination paired with corresponding short exposure integration times. Spectrally, the infrared emitting diodes were matched to narrow-band spectral filters on the cameras.

Another challenge is acquiring video of the insect, rather than a recording of a bright point of light on a dark background. To overcome this challenge, we used a selective transmission mirror to direct the infrared light path to the tracking camera and visible light to a high-speed video camera which could be triggered to record at 1000 or more frames per second. Obtaining sharp, well-illuminated videos with this system, however, requires the use of a long focal length objective with a large aperture and, consequently, a large diameter optical path which is redirected to follow the insect. Therefore, we developed a large size pan-tilt mirror periscope turned by stepper motors with a design goal of minimal mass, and therefore minimal inertia and thus latency.

A large aperture, long focal length objective has a shallow depth of field, and thus a final challenge is maintaining focus on a flying insect. We used a second tracking camera to implement a stereo camera pair. Distance estimates from stereo disparity get less precise with increasing distance, but this decreased distance precision is matched by an increased depth of field at larger distances.

We combined these design elements in a single system capable of recording high-speed, high-magnification video of freely flying insects in natural, outdoor settings (Fig. 3).

**Fig. 3.**
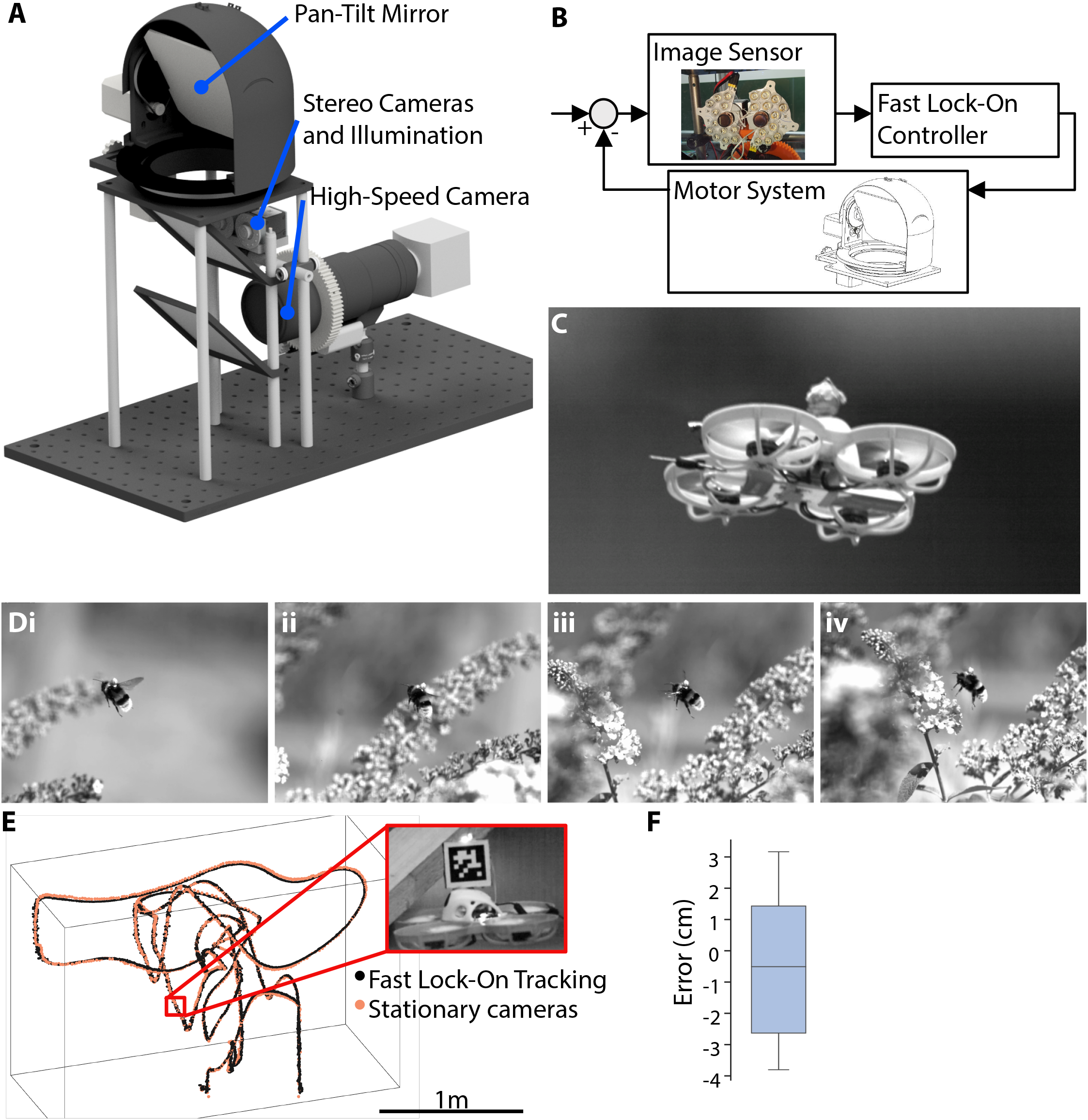
Large diameter optical path shared with high-speed telescopic video camera to record high-resolution insect video during natural behavior. (A)A large diameter optical path is be steered by a Fast Lock-On system incorporating active infrared illumination, and a low-latency infrared stereo camera pair. A wavelength-selective mirror allows high-speed videography of the scene along the same optical axis. Stereo disparity is used to control a focus motor on a high-magnification objective. (B)The Fast Lock-On core uses one camera from a stereo camera pair and paraxial infrared illumination to provide feedback about the position of a small retroreflective marker which is then used to drive a motor pair to re-orient the optical axis over a large angular range. (**C**) Example image from a high-speed, high-resolution video of a toy quadcopter. (**D**) Example frames from a high-speed video of a bumble bee landing. Marker size: 3 mm diameter, mass: 20 mg. (**E**) Simultaneous three-dimensional tracking with a conventional motion capture system using five high-resolution stationary cameras (orange points) and a single FLO system (black points) over several meters. Inset shows single frame from a 300 frame per second high-speed video acquired using high-speed camera directed with FLO system. (**F**) Absolute tracking errors quantified by measuring 1500 millimeter length rod in 12 configurations distributed throughout a multiple meter tracking volume. Box shows quartiles with median show as line and whiskers show entire data range. N = 88. **Movie 1. High-speed video of bumble bee captured using fast lock-on tracking**.

To validate the FLO based high-speed, high-resolution video system, we tracked a toy quadcopter to which a retroreflective marker was attached. Testing confirmed all system components functioned and that high resolution, high-speed videos could be acquired over a large field of view and the stereo-based autofocus system in an indoor setting (Fig. 3C, Movie S1).

We next moved our system outdoors with the goal of recording high-speed, high-resolution video of insects behaving in naturalistic conditions. After marking insects with retroreflective markers, we were indeed able to track them with our FLO system and record videos in which the head-tail length of the insect is hundreds of pixels long with little or no motion blur (Fig. 3D, Movie 1, S2, S3, S4). The mass of the reflective marker ball attached to the bees was about 20 mg. For bumble bees *Bombus terrestris*, this compares to an average forage load of 47 mg [26], and in *Apis terrestris* of 17-25 mg [27]. We did not observe any obvious changes in behavior or other effects due to the added mass, aerodynamic changes, or other perturbations caused by our markers.

### Accuracy and precision of FLO-based position measurements

To obtain an estimate of the performance of this system in producing three-dimensional position estimates of the tracked object, we returned indoors where we acquired a dataset in which the toy quadcopter was tracked with both the FLO system and a conventional motion capture system using stationary cameras (Movie S5). In the FLO-based system, the pan and tilt motor positions, together with the disparity measurement from the stereo tracking cameras, were used with to compute spherical coordinates which were then converted to cartesian coordinates. We flew the quadcopter though a space filling an axes-aligned bounding box of 2.3 meters by 1.0 meters by 1.5 meters. The FLO-based system and the stationary camera system showed close agreement in estimated 3D coordinates (Fig. 3E). To obtain a quantitative estimate of absolute tracking accuracy and precision with this FLO system, we affixed retroreflective markers to the ends of a 1500 millimeter rod (measured with a 5 meter tape measure from Duratool) and performed 88 measurements of its length throughout a multiple-meter volume. The rod was held in multiple configurations such as oriented almost directly away from the FLO system, oriented perpendicularly to the line of sight from the FLO system, and at a variety of positions in the volume. The mean absolute error was 1.96 centimeters, or 1.3%, and the distribution of errors is shown in Fig. 3F.

### Recording bee trajectories from a flying quadcopter drone

Finally, we sought to test the feasibility of tracking bees from a flying quadcopter using a lightweight, self-contained FLO system (Fig. 4A). For this system, the image sensor and illumination source were the same as described above used to record high-speed, high-resolution videos (Fig. 4B). With this system, we were indeed able to track marked honey bees for several minutes and could measure the pan tilt and distance from the drone to the tracked bee. A drone pilot followed the bee with the drone to keep the bee within tracking range of the drone. Data fusion with GPS and attitude data from the drone allowed reconstructing the 3D trajectory of the bee (Fig. 4C,D,E, Movie 2). With the present system, we anticipate that tracking will be limited to areas where no specular reflections of the LED other than the insect-mounted retroreflector are present.

**Fig. 4.**
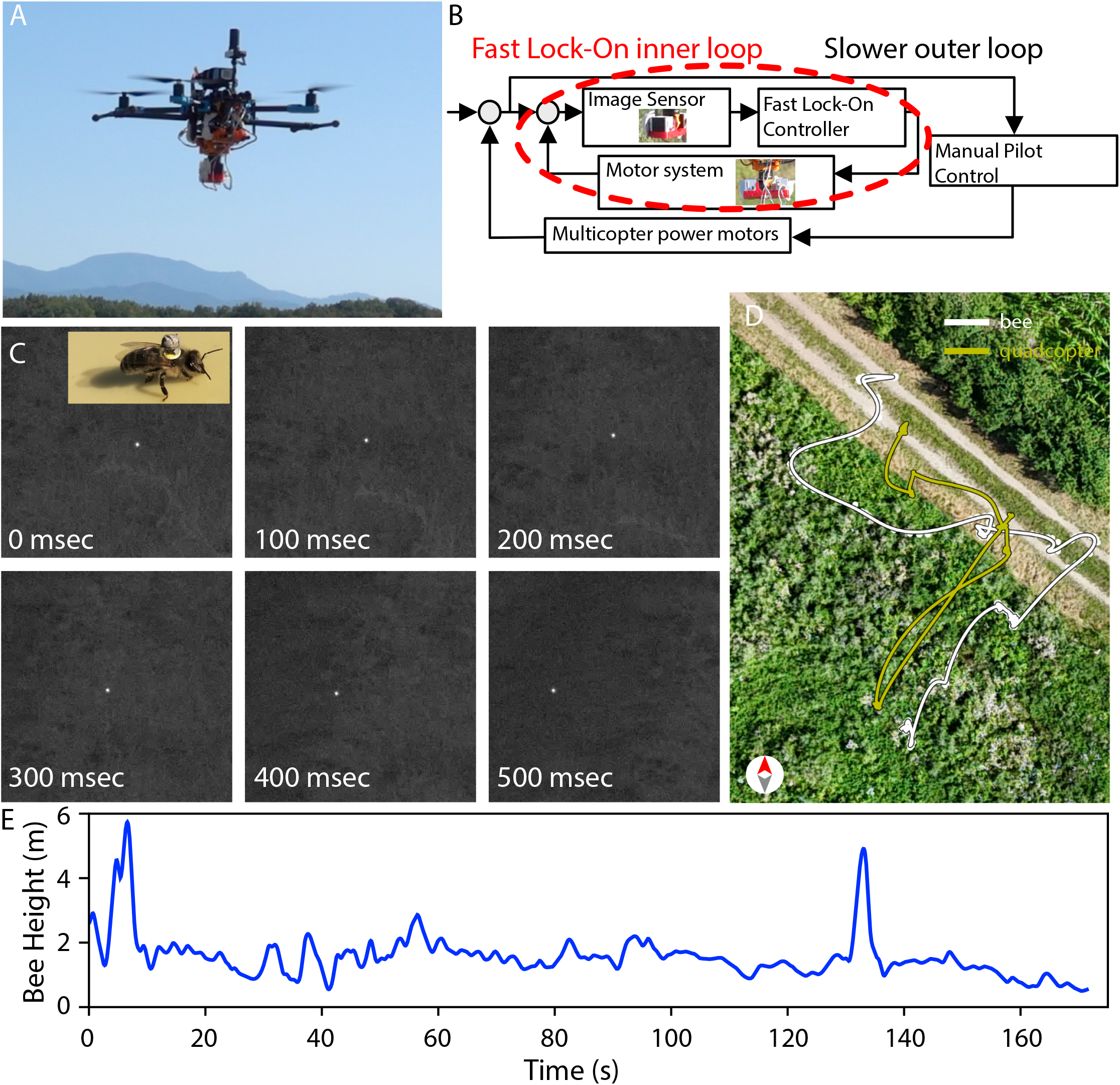
Quadcopter mounted fast lock-on system for tracking insects. (**A**) A quadcopter is equipped with a fast lock-on system optimized for low mass. (**B**) The sensor and illumination is a stereo pair of video cameras with paraxial infrared illumination. Images are processed on a PC carried onboard the quadcopter and drive the pan-tilt motor system automatically. Bee angular position information is conveyed to the pilot who attempts to steer the drone such that it remains within tracking range of the bee. (**C**) Example video frames acquired during tracking a marked honey bee *Apis mellifera*. (**D**) map of tracked 3D bee position plotted over aerial photo of location (**E**) height of bee computed during flight shown in D. **Movie 2. Tracking a bee with a quadcopter mounted fast lock-on system**.

## DISCUSSION

Here we described Fast Lock-On (FLO) tracking and used it to acquire high-speed, high-resolution videos of insects flying outdoors. These videos filmed the insect from takeoff to landing with high magnification and low motion blur. Appendages such as legs, wings, and antennae remained in focus, even as the insects flew. We further demonstrated a FLO system mounted on a quadcopter to tracking the position of flying honey bees.

As a methodology, Fast Lock-On tracking has some drawbacks. Closed-loop tracking requires that an initial lock must be established via a separate process and, if tracking is lost momentarily, this may result in a complete loss of tracking. Another drawback is that distance estimation must be performed additionally to the basic fast lock-on concept. Here we partially overcame this problem with stereopsis to measure distance and adjust focus. Other solutions, such as measuring time of flight of light pulses, are worth investigating as alternatives. Furthermore, while successful tracking does not require careful calibration, extracting final 3D coordinates and estimating their accuracy will require careful calibration.

Computational demands from image processing were minimized by the use of a retroreflective marker in the systems described here. Affixing the mark to an insect, however, requires physically manipulating the insect to be tracked and, although we did not observe any obvious example, it potentially causes interference with behavior. The requirement to use retroreflectors could be lifted if more sophisticated computer vision methods could be developed to perform the require task. It would be worthwhile to investigate the use of marker-free approaches making use of object detectors based on artificial neural networks such as YOLO [28] or event cameras [29].

Several further improvements are conceivable. The use of multiple FLO systems working together would theoretically allow calculation of 3D coordinates and distance estimation via triangulation. These systems could operate independently and the data combined afterwards for processing, but if they would work cooperatively during tracking, improved robustness to occlusions and larger tracking volume coverage could also be acheived. This would involve calibrating multiple systems in a common frame and implementing higher-level coordination such as sending, receiving, and using estimated positions between individual FLO systems. For the drone-based systems, this could involve using real-time kinematic global navigation satellite system receivers in conjunction with onboard inertial measurement units.

We suggest FLO tracking could potentially be applied to study the biomechanics of insect landing behavior, to study the orientation of the head and eyes during learning flights of bees, to study at what altitude bees fly, to study responses of insects to artificial light at night, to pesticide treatment, or to habitat loss. All of these research topics have open questions and progress is presently limited by the technology available for recording insect behavior.

## MATERIALS AND METHODS

### Fast Lock-On controller

The closed-loop controller is a standalone executable written in the Rust programming language which can be compiled for Linux, Windows and macOS operating systems. The core inner loop updates two Kalman filters where the prediction step consists of computing the predicted (a priori) state estimate and covariance. The first Kalman filter estimates the four-dimensional state 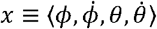 representing pan angle, pan angular velocity, tilt angle, and tilt angular velocity in a coordinate frame fixed relative to the base of the fast mirrors (i.e., stationary in the world frame except in the case of the drone). The second Kalman filter estimates the two-dimensional state 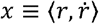 of distance and rate of change of distance. Observations for the pan-tilt system are the sensor coordinates provided by the Imops module of Strand Camera. Ongoing lag-compensated estimates of motor angles are maintained via non-probabilistic representations empirically fit to motor data. Motor commands are then computed each cycle through the loop. A standard PC was used to run the controller software. In the case of the drone, to reduce mass, this PC (ASRock 4×4 Box-5800U) was mounted to the drone without its housing.

### Angle transformations

In order to provide the Kalman filter with observations of the target position in global ‘motor angle’ coordinates *ϕ*_*t*_ (pan) and *θ*_*t*_ (tilt), these are reconstructed form the current (estimated) motor position *ϕ*_0_, *θ*_0_ and the apparent target position in ‘sensor angle’ coordinates *x* and *y*, using the linear approximation

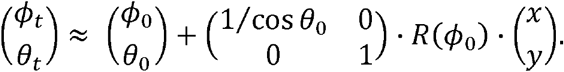

Depending on the specific system geometry, the image does (‘moving mirror’ setup) or does not (‘moving camera’ drone setup) pick up a roll angle equivalent to *ϕ*_0_, which is reflected by different choices

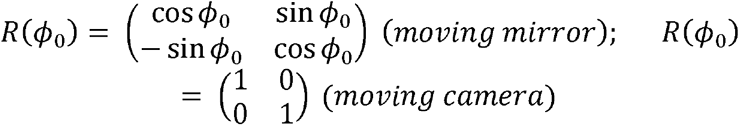

for the two setups.

### Cameras, image acquisition, and infrared illumination

Cameras used for low latency image acquisition were a digital camera capable of 160 uncompressed frames per second in full frame mode and more than 1000 frames per second with a small region of interest (Basler ace 2 a2A1920-160umBAS). For experiments involving stereo images for distance estimation, an external trigger device (https://github.com/strawlab/triggerbox) provided voltage pulses to synchronize acquisition. Image acquisition and low-latency processing was performed by Strand Camera 0.12 (Straw Lab, https://strawlab.org/strand-cam). Image processing was implemented in the Imops module of Strand Camera, which implements SIMD-accelerated operations to find the location of the brightest pixel. For experiments with the high-speed video camera and the drone, a custom circuit board carrying high intensity infrared LEDs (SFH 4715AS A01, Osram) was used as an illumination source and Thorlabs FBH05850-10 bandpass filters were used to cut other sources of illumination. For all systems, a 25mm focal length objective (IDS-5M12-S2524F, IDS Imaging) was used.

For high-speed, high-resolution video, we used a long focal length macro lens (Nikon AF Micro Nikkor 200mm f/4D) and camera (Mikrotron MotionBlitz EoSens mini1). The motion capture system used to obtain independent 3D measurements of the quadcopter trajectory was Braid 0.12 (Straw Lab, https://strawlab.org/braid) using 5 Basler ace 2 a2A1920-51gcBAS cameras.

### Design considerations for active illumination and retroreflectors

The principal requirement is that the brightness of the retroreflector as seen by the camera (pixel value) is higher than the brightest parts of a sunlit environment. If the image of the reflector is larger than one pixel in width, the pixel value is approximated by:

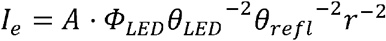

*I*_*e*_ is the illuminance of the sensor (pixel value), *Φ*_*LED*_ is the output light flux of the led, *θ*_*LED*_ is the angular divergence of the LED beam, *θ*_*refl*_ is the angular divergence introduced by the retroreflector, r is the distance to the marker, and the factor A captures the contributions of everything else (constants and unit conversions, losses in LED reflector lens, lens aperture, filter transmission, retroreflector loss).

(assumptions: camera lens is focused on the marker; *θ*_*LED*_, *θ*_*refl*_ << 1; lens pupil < *r · θ*_*refl*_; LED to pupil distance < *r · θ*_*refl*_)

The size of the spot decreases as 1/r as the marker is moved further away from the camera, but that is not an issue until the size gets smaller than one pixel (or the diffraction limit of the lens). Beyond that, an additional *r*^-2^ multiplier will creep in, i.e. the pixel value will now be determined by the total energy received by the camera, not the illuminance at the center of the spot. This falloff (*r*^-4^) is extremely fast, so the point where the reflector size corresponds to one pixel of the camera can be considered a hard limit for how far the tracking can work.

We have experimentally verified that pixel values do indeed fall off as *r*^-2^ when the image of the reflector is much larger than 1 pixel.

It may be useful to consider how to double the tracking distance. First, *θ*_*LED*_ should be quadrupled. Then, to avoid the *r*^-4^ falloff, either the magnification of the lens could be increased to fill more pixels with the image of the reflector, the size of the reflector could be increased to fill more pixels, or an intermediate combination of the two. As the image of the reflector scales linearly with the reflector radius, the reflector linear dimension should be doubled if the lens magnification was maintained. We note that increasing the magnification may decrease tracking robustness as it may also result in slower movement and decreased field of view. When considering miniaturizing the reflector, the same considerations apply and one should choose system components such that the reflector image remains larger than one pixel.

### Motor systems

The mini-FLO system uses hobby PWM servos arranged in a pan-tilt configuration (RB-Lyn-74, Robotshop) onto which the camera physically mounted. The high-resolution camera system uses smart stepper motors (PD42-2-1240-TMCL, Trinamic). The drone uses brushless gimbal motors (GM2804H-100T Encoder Combo Set, iFlight). The drone itself was a Holybro X500 V2 with Holybro H-RTK F9P Helical GPS RTK Module.

### Software and design files

Upon acceptance for publication, the source code for the Fast Lock-On controller and the design files tracking systems will be released under an open-source license. The other software components developed are already under an open-source license.

### Reflector marker attachment to desert locust

Winged adult stage desert locusts, *Schistocerca gregaria*, were obtained from BUGS-international GmbH (Irsingen, Germany). Locusts were housed in groups of up to 8 within a vivarium (30×30×40cm) in laboratories, and fed wet green lettuce daily, until individuals were removed for testing.

Markers attached to locusts were made of several sections of retroreflective tape (OpitTrack 3M 7610 reflective material, NaturalPoint Inc., Corvallis, Oregon, USA) and adhered to the insect using the tapes’ own adhesive backing (see Fig. 1D). First, a ‘saddle’ or trapezoid shaped section was applied across the pronotum section of the locust’s thorax (approx. 20mm and 10mm wide at the long and short end, and 10mm deep). Care was taken to not apply this section of tape beyond the pronotum. Second, a rectangular section was applied to the front of the insects’ head (approx. 2mm by 5mm), avoiding the antennae and compound eyes. Third, a rectangular section was applied to ventral surface of the locust’s thorax (approx. 6mm by 6mm), with care taken to not restrict movement of the locust’s legs. In some trials (as seen in Fig. 1D) a fourth section was applied to the ventral surface of the locust abdomen (approx. 6mm by 6mm); however, this forth section was rarely visible while other sections were not and was therefore not applied in later trials.

All sections were cut to size after direct comparison with the insect to be marked. Thus, marker sections varied in exact size and shape (note all sizes reported above are approximations). Locusts were not anesthetized during attachment of markers. Instead, markers were attached while the insect’s hind legs were held across the femurs and abdomen, preventing escape behavior. All sections were applied with nitrile-gloved hands and pressed down further with a blunt-seeker tool. If the marker sections did not adhere easily to the hard sections of the insects’ bodies, these sections could be lightly scraped with an emery file to rough up this surface and facilitate tape adhesion, however this was rarely required. Collectively the marker sections applied to the locust weighed on average (mean) 70mg (based on 5 measurements of tape sections ranging between 64mg and 76mg).

### Reflector marker attachment to bees

Bumble bee colonies, *Bombus terrestris*, were obtained from colonies purchased from Biobest (Westerlo, Belgium), while honey bees, *Apis mellifera*, were obtained from hives maintained on-site. Honey bee colonies were maintained by standard beekeeping practices by an on-staff beekeeper, however colonies were kept outside and bees were otherwise required to forage on local flora for food. As these bees were raised commercially, no permit for working on wild insects is required.

The markers attached to both bee species were of identical design and procedures for attachment of markers were also identical for both species. Bee retroreflective markers consisted of a 3mm retroreflective sphere attached to a queen marking plate (Opaliths, Pasieka, Dolna, Poland) stuck to the bee’s back (Fig. 1D). Retroreflective spheres were made as follows. Small 3mm polystyrene balls (TEDi GmbH & Co. KG, Dortmund Germany) were wrapped in a 20mm by 3mm strip of retroreflective tape, stretched as applied to minimize creasing, until the complete surface of the ball was covered. This retroreflective sphere was then stuck directly to the outward face of a queen (bee) marking plate with super glue (Loctite original, Henkel Corporation, Westlake Ohio, USA). Use of the marking plate as a base of the marker allowed the sphere to be held higher and allowed better and cleaner attachment of the marker to the bee, compared to directly sticking the sphere to the bee. Such plates fit the bee thorax and limit spread of glue applied beneath them. Markers were prepared, as described above, before bee capture to minimize time bees spent captive.

Foraging bees from colonies were caught using butterfly nets upon exiting the colony. Bees were not anesthetized for marker attachment, instead captured bees were transferred to a ‘queen marking’ or ‘drawing tubes’ with net head and foam slider to hold bees in place (Josef Muhr Beekeeping, Prakenbach, Germany). In these bees were positioned and held still against a net grid with the dorsal section of their thorax exposed through the net grid. The dorsal section of the thorax was shaved of hair with a razorblade, doing so allows marker plates to be stuck directly to the bee’s cuticle not its hair, further facilitating adhesion. The marker was then stuck to the bee using the marking plate base (Fig. 1D) as the point of contact using a small amount of (Loctite) super glue applied to the bee with a pipette tip. As when using normal queen marking plates, the marker was positioned in the center front of the thorax and care was taken to not apply glue to or otherwise obstruct the bee’s wings. Marked bees were then transported to falcon tubes (with air holes) for release. Bee markers weighed 20mg (mean average of 18 markers, ranging between 16mg and 28mg).

## Supplementary Materials

Movie S1. High-speed video of quadcopter

Movie S2. High-speed video of locust

Movie S3. High-speed video of honey bee

Movie S4. High-speed video of bumble bee

Movie S5. Simultaneous high-speed video of quadcopter and stationary camera system Supplementary ZIP file containing design files, build and usage instructions, and source code. This material will be publicly released in the open-source GitHub repository https://github.com/strawlab/flo upon publication.

## Acknowledgments

We thank Mathias Siegel and the mechanical workshop of the Institute of Biology I for constant assistance, Matthias Wittlinger for use of the high speed video camera, Julius Klein for assistance with bees, and Erwin Wagner for use of the field used in the drone experiments.

## Author contributions

Conceptualization: TTVD, ADS

Funding acquisition: TTVD, ADS

Investigation: TTVD, VVT, MJMH, SL, ADS

Methodology: TTVD, VVT, MJMH, SL, ADS

Project administration: ADS

Software: VVT, SL, ADS

Supervision: ADS

Visualization: TTVD, ADS

Writing – original draft: ADS

Writing – review & editing: TTVD, VVT, MJMH, SL, ADS

## Data and materials availability

All data are available in the main text or the supplementary materials. Supplementary ZIP file contains design files, build and usage instructions, and source code. This material will be publicly released in the open-source GitHub repository https://github.com/strawlab/flo upon publication.

